# RNA-dependent RNA amplification as a strategy to perform tRNA sequencing on low-input samples

**DOI:** 10.64898/2026.08.24.746598

**Authors:** Élie M. Teyssonnière, Mari Mito, Yuichi Shichino, Shintaro Iwasaki

## Abstract

Transfer RNAs (tRNAs) are key molecules that deliver amino acids to the translating ribosome according to their cognate codon encoded in messenger RNA (mRNA). Due to the modified nature of tRNA nucleotides, accurate tRNA quantification can be tedious, especially when dealing with small sample inputs. Here, we took advantage of an RNA-dependent RNA amplification method using T7 polymerase to quantify tRNA abundance in low biological input. Our method, called T7 High-resolution original RNA (Thor)-tRNA-Seq, showed reproducible and quantitative measurement of low tRNA inputs. Thus, our Thor-tRNA-Seq is a robust and reliable approach for the quantification of tRNA in samples with small and precious biological material.

## Introduction

Translation of mRNA sequences into proteins requires tRNA as an amino acid carrier and codon decoder for peptide synthesis. In the cell, the pool of tRNA available is a critical factor impacting translation regulation ^1–5^. For instance, elongation speed can be drastically modulated upon changes in tRNA abundance ^6–11^, and such a phenomenon results in a strong impact on protein evolution, with protein sequence being adjusted to match the number of tRNA gene copies in the genome ^12,13^. Improper tRNA abundance or regulation is a known source of cellular dysfunction and has been linked to several human diseases ^14–16^. In this context, quantifying the pool of cellular tRNA has played a central role in understanding the effect of abnormal tRNA abundance ^17^. In recent decades, along with the development of next-generation deep sequencing techniques, such quantification has been greatly facilitated ^18^.

However, tRNA sequencing (tRNA-Seq) requires significant adaptation of the conventional RNA sequencing (RNA-Seq) protocol, because of the short length, amino acid conjugation at the 3′ end, and high rate of RNA modification ^19^. To overcome these inherent challenges of tRNAs, diverse derivatives have been developed that incorporate specialized linker-ligation strategies, enzymatic removal of RNA modifications, and modification-resistant reverse transcriptases ^20^. However, one remaining limitation of these approaches is the fact that they rely on a consequent input of biological material and can be limited when exploring subtle tRNA amounts or small changes in tRNA content ^20^. This can be problematic, especially with the recent direction of biological exploration aiming toward more precise quantification using small biological input ^21,22^, subcellular explorations ^23^, or co-purification-based sampling ^24,25^.

In this work, we developed low-input tailored tRNA-Seq, which harnesses RNA-dependent RNA amplification before sample loss during library preparation. The T7 High-resolution original RNA (Thor) technique uses T7 RNA polymerase ^26^, which can transcribe RNAs complementary to DNA linker-ligated RNAs. Indeed, the introduction of Thor to ribosome profiling or Ribo-Seq allowed robust, reproducible, less biased libraries from ultra-low materials ^27^. The Thor approach enables us to generate antisense RNAs of highly modified tRNAs and stable library preparation. Overall, our results support that Thor-tRNA-Seq is a powerful approach to reliably explore low-input tRNA samples.

## Results and Discussion

### Amplification of tRNA by T7 RNA polymerase

Sequencing the tRNA pool in cells can suffer from limitations, especially when dealing with samples with low amounts of biological material. To overcome this problem, we took advantage of a recently developed technique, Thor, which relies on RNA-templated *in vitro* transcription of RNA fragments using T7 RNA polymerase (Figure 1A, top) ^27^. Ligation of the DNA linker containing the antisense sequence of the T7 promoter, followed by hybridization with a complementary DNA oligonucleotide, generates a partial dsDNA promoter region, enabling RNA-templated transcription of the downstream sequence ^26,28,29^. This strategy benefits from 1) the material multiplication at the early step of library construction before sample loss and 2) the linear amplification that is less biased than exponential amplification in PCR ^30^. Although originally this method was developed to handle short ribosome-protected RNA fragments (or ribosome footprints) in Ribo-Seq ^27^, we sought to adapt this technique and see if it could be used for tRNA, where strong RNA structure and RNA modification are known to affect sequencing experiments.

**Figure 1.**
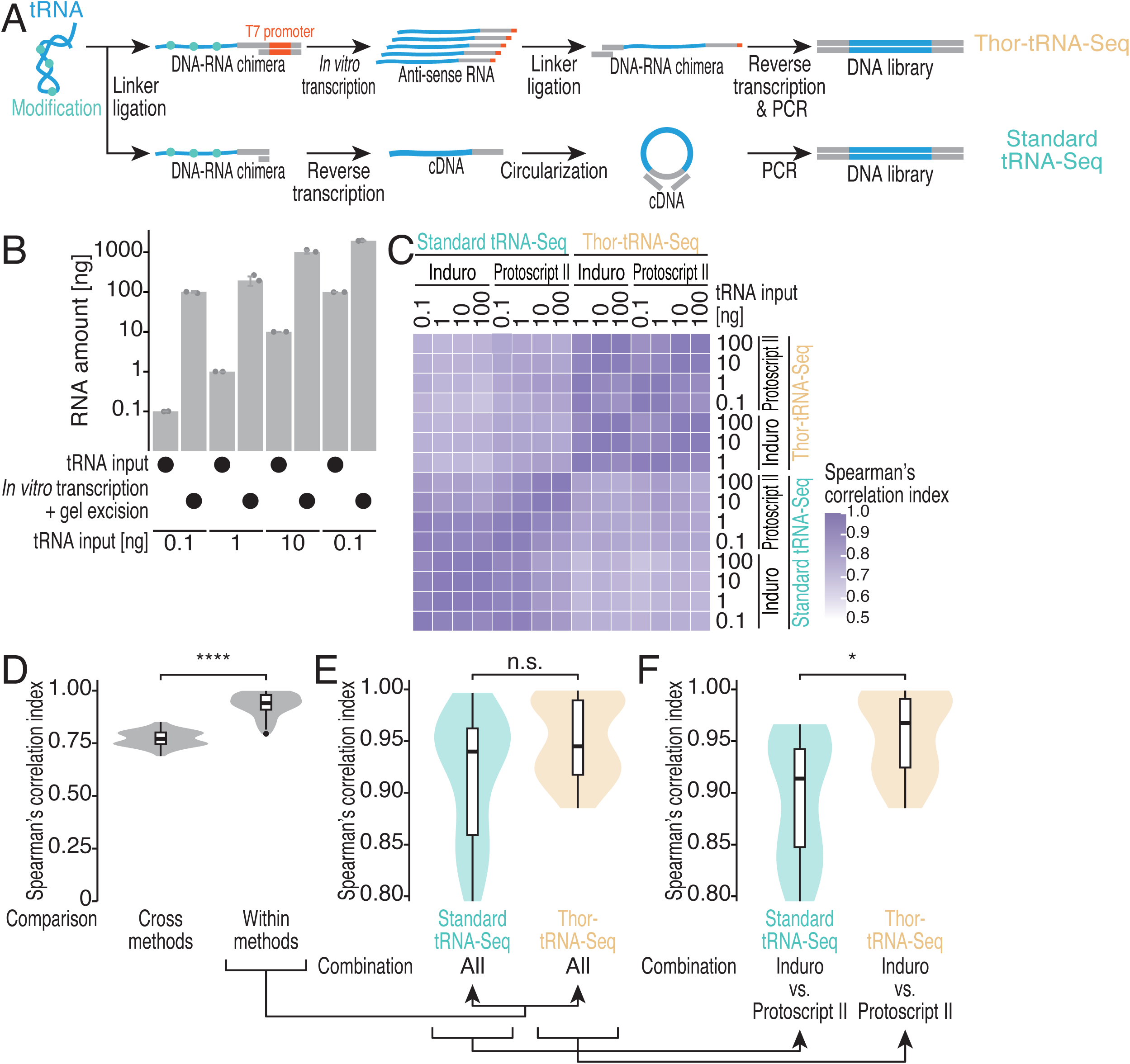
Robust and comprehensive quantification of low-input tRNAs by Thor-tRNA-Seq. (A) Schematic workflow of Thor-tRNA-Seq and standard tRNA-Seq. (B) The amount of input RNA and amplified RNA after *in vitro* transcription for the indicated conditions. The data are presented as the means (bars) for replicates (points, n = 2). (C) Heatmap of pairwise Spearman’s correlation coefficients between Thor-tRNA-Seq and standard tRNA-Seq datasets from independent replicates under indicated conditions. The correlation coefficients are represented by the color scale. (D) Box and violin plots of correlation levels across methods and within methods. (E) Box and violin plots of correlation levels within standard tRNA-Seq and Thor-tRNA-Seq. (F) Box and violin plots of correlation levels comparing Induro and Protoscript II reverse transcriptases within standard tRNA-Seq and Thor-tRNA-Seq. Box plots show the median (centerline), upper/lower quartiles (box limits), and 1.5× interquartile range (whiskers). Significance was determined by the Mann‒Whitney *U* test. ***, *P* value < 0.001; **, *P* value < 0.01; *, *P* value < 0.05 See also Figure S1.

To test the potential of the material amplification by *in vitro* transcription, we titrated amounts of purified yeast tRNA input: 100, 10, 1, and 0.1 ng. While no linker-ligated tRNA material (∼150 nt in total) was visible for 1 and 0.1 ng input by nucleic acid staining on the gel (Figure S1A), the antisense RNA products appeared after *in vitro* transcription (Figure S1B). For all the input conditions, we observed an increase in material by *in vitro* transcription (Figure 1B), ranging from 95- to 1970-fold changes (Figure S1C); the lower input resulted in the highest tRNA increase. The antisense RNAs of tRNAs were used in the downstream library preparation (Figure 1A, top). Although tRNA modifications pose a barrier to readthrough by reverse transcriptase ^31–34^, *in vitro* transcription by T7 polymerase may overcome this challenge, magnifying the materials.

For reference, we compared our Thor-tRNA-Seq approach with a conventional tRNA-Seq method, which employs direct reverse transcription of linker-ligated tRNA to convert cDNA (Figure 1A, bottom). As the choice of reverse transcriptase influences the extension block and the nucleotide misincorporation of reverse transcription by modified bases in tRNA, we tested two different enzymes that have been used for tRNA-Seq: Protoscript II ^35^ and Induro ^32^, applying a long incubation (16 h) strategy as conducted in modification-induced misincorporation (mim) tRNA-Seq ^31^ to maximize the yields of the full-length cDNA (Figure S1D-E). These standard tRNA-Seq experiments, with 10 ng or less, were technically challenging since the cDNA was barely visible on the gel (Figure S1D-E). This highlighted the advantage of nucleic acid amplification with the Thor technique (Figures 1B and S1B). In standard tRNA-Seq, we pooled the barcoded cDNAs from all the titrated inputs for the downstream process, allowing the 100 ng input sample to act as a carrier for the other low-input samples.

The two reverse transcriptases were also used for Thor-tRNA-Seq (Figure S1F-G), although RNA modifications should be erased in the *in vitro* transcribed antisense RNA of the tRNA (Figure 1A, top; note that a 0.1 ng input sample with Induro was omitted due to a quality issue).

### Robust quantification of low-input tRNA by Thor-tRNA-Seq

Next, we sought to compare the quantitative output of the standard tRNA-Seq and Thor-tRNA-Seq. Counting the reads mapped to the tRNA sequence reference of *Saccharomyces cerevisiae* ^36^, we computed Spearman correlation indices (ρ) for each method comparison across titrated inputs (Figure 1C). Within-method comparison (*i.e.*, correlation among standard tRNA-Seq samples or correlation among Thor-tRNA-Seq samples) displayed higher correspondence than cross-method comparison (*i.e.*, correlation between standard tRNA-Seq and Thor-tRNA-Seq) (Figure 1D). We found that Thor-tRNA-Seq showed higher within-method correlation than standard tRNA-Seq (Figure 1E), although the difference did not pass the statistical threshold. The variance in standard tRNA-Seq stemmed from the differences in reverse transcriptase (Figure 1F; comparisons restricted to cross reverse transcriptases). In contrast, Thor-tRNA-Seq showed higher consistency among the samples, irrespective of reverse transcriptases (Figure 1E-1F); this was due to the unmodified RNA templates for reverse transcriptase in Thor-tRNA-Seq (Figure 1A). Our data showed high reproducibility in Thor-tRNA-Seq, providing a correlation coefficient ranging between 0.885 and 0.999, with a median of 0.940 (Figure 1E). Our Thor-tRNA-Seq also presented relatively better correspondence with earlier tRNA-Seq work ^37^ (Figure S1H).

### Potential application of low-input-tailored Thor-tRNA-Seq

Adequate methods to deal with small amounts of biological material are of central need for experimental biology, as major techniques such as single-cell analysis, subcellular exploration, or co-precipitation result in small biological input for downstream experiments. Such a method was, to our knowledge, nonexistent for tRNA-Seq. In this work, we took advantage of a recently developed technique, Thor-Ribo-Seq ^27^, that increases small biological mRNA samples using *in vitro* transcription and checked whether this could be used in the context of tRNAs. Our resulting method, Thor-tRNA-Seq, showed an overall good agreement with the conventional tRNA-Seq approach even with low inputs. The same approach should be applicable to other short non-coding RNAs, including small interfering RNAs (siRNAs), microRNAs (miRNAs), PIWI-interacting RNAs (piRNAs), and small nucleolar RNAs (snoRNAs).

## Methods

### tRNA processing

Several input quantities (100, 10, 1, or 0.1 ng) of commercially available yeast tRNA (AM7119, Thermo Fisher Scientific) were dissolved in 0.1 M Tris-HCl, pH 9.0, and deacylated at 37°C for 45 min. The tRNA was purified with an Oligo Clean & Concentrator Kit (D4060, Zymo Research), denatured at 95°C for 2 min, and then dephosphorylated using a T4 polynucleotide kinase (M0201S, New England BioLabs [NEB]) as described previously ^38^.

### Standard tRNA-Seq

Pre-adenylated linkers (NI-810 to NI-817) were prepared with a DNA Adenylation Kit (E2610L, NEB) and ligated to the processed tRNA (see “tRNA processing” section) with T4 RNA Ligase 2, truncated KQ (M0373L, NEB), as described previously ^38^. The samples were then cleaned using an Oligo Clean & Concentrator Kit (D4060, Zymo Research) and loaded onto a 15% denaturing polyacrylamide gel. After separation by electrophoresis, tRNA ligated with the linkers (approximately 130 nt long) was excised from the gel and purified as described previously ^38^. The purified linker-ligated tRNA was hybridized with a reverse transcription primer (NI-802) ^38^ by incubation at 65°C for 5 min and cooling on ice. The reverse transcription was performed with Protoscript II (M0368L, NEB) for 16 h at 49°C ^35^ or Induro (M0681, NEB) for 16 h at 42°C ^32^. Then, tRNAs were hydrolyzed using 0.1 M NaOH at 70°C for 20 min. After another round of purification with an Oligo Clean & Concentrator Kit, the samples were run on a 10% denaturing polyacrylamide gel, and the cDNAs were excised from the gel and purified. The recovered cDNAs were pooled (8 samples; 4 samples with Protoscript II and 4 samples with Induro) and circularized by CircLigase II (CL9025K, LGC, Biosearch Technologies) as described previously ^38^. The circularized cDNAs were used for PCR amplification, run on a 15% non-denaturing polyacrylamide gel, and excised as described previously ^38^. Ultimately, 10 PCR cycles were required to recover enough DNA for sequencing.

### Thor-tRNA-Seq

We adapted the Thor-Ribo-Seq technique ^27^ to perform tRNA-Seq. In brief, pre-adenylated linkers (SI96_089 to SI96_096) ^27^ were ligated to the processed tRNA (see “tRNA processing” section) with T4 RNA Ligase 2, truncated KQ (M0373L, NEB). After column purification with an Oligo Clean & Concentrator Kit (D4060, Zymo Research), the ligated tRNAs were also run on a 15% denaturing polyacrylamide gel and excised. The resulting linker-ligated tRNAs (approximately 150 nt long) were then used for *in vitro* transcription with a T7-Scribe Standard RNA IVT Kit (C-AS3107, CELLSCRIPT), incubating the reaction for 2.5 h at 37°C. The reaction was then treated with DNase I for 20 min at 37°C. The samples were cleaned and purified using RNA Clean XP beads (A63987, Beckman Coulter). After measuring concentration using a Qubit RNA HS Assay Kit (Q32852, Thermo Fisher Scientific), 300 ng of RNA was loaded and run on a 10% denaturing polyacrylamide gel. If the *in vitro* transcription of low tRNA inputs resulted in smaller RNA amounts (175.4 ng and 91.8 ng for the 10 and 0.1 ng input, respectively), the entire tRNA quantity was run and used for the later procedure. The amplified tRNAs, after being excised from the gel and purified, were ligated with a second linker (SI-183) with T4 RNA Ligase 2, truncated KQ (M0373L, NEB) ^27^. At this step, the sample for Induro reverse transcriptase with 0.1 ng tRNA input was lost because of DNA contamination. The 2nd linker-ligated RNAs were then used for reverse transcription with Protoscript II and Induro. The pooled samples (2 pools [Protoscript II pool and Induro pool] with 4 samples each) were also amplified with PCR, and 6 PCR cycles were required to obtain a sufficient amount of DNA for sequencing.

### Sequencing and data processing

The libraries were sequenced using Illumina sequencing on a NovaSeq X Plus platform, using a paired-end (PE) 150-bp option. The resulting reads were then trimmed of their linker using fastp (version 0.21.0) ^39^ and split using cutadapt (version 3.7) ^39^. UMI in the reads was retrieved using UMI-tools (version 1.1.4) ^40^. The resulting reads were mapped to the tRNA genes from the *S. cerevisiae* reference genome ^36^, with the addition of CCA at the end of each tRNA sequence. The mapping was done using STAR (version 2.7.0a)^41^. We extracted the mapped reads and indexed them using SAMtools (version 1.10) ^42^. Then, the duplicates generated in *in vitro* transcription (3′ UMI) (Thor-tRNA-Seq) or PCR (both 5′ and 3′ UMI) (Thor-tRNA-Seq and standard tRNA-seq) were removed using UMI-tools, and the mapped read count for each tRNA was extracted.

### Data analysis

The data analyses were all conducted using R (version 4.4.2). Correlation between the raw counts + 1 obtained for each method was computed using the Spearman correlation coefficient. In all later analyses, differences between groups were always computed using Wilcoxon tests, unless explicitly specified. The tRNA sequence coverages and mismatches were obtained using the Rsamtools package (version 2.22.0) ^43^. The tRNA intron sequences were defined using the *Saccharomyces* Genome Database (SGD) (https://www.yeastgenome.org) and removed from the analysis when specified.

## Acknowledgements

We are grateful to all the members of the Iwasaki laboratory for their technical help and constructive discussions. For this work, we employed the HOKUSAI SailingShip supercomputer facility at RIKEN. This work was supported by the Ministry of Education, Culture, Sports, Science and Technology (MEXT) (JP24H02307 to S.I.; JP21H05734, JP23H04268, and JP25H01440 to Y.S.), the Japan Agency for Medical Research and Development (AMED) (JP20gm1410001 to S.I.; JP23gm6910005 to Y.S.), the Japan Society for the Promotion of Science (JSPS) (JP25KF0086 to E.M.T.; JP23H00095 and JP26H02372 to S.I.; JP21K15023, JP23K05648, JP26K01935, and JP26K23132 to Y.S.), the Japan Science and Technology Agency (JST) (JPMJCR25T2 to S.I. and Y.S.), the Nakajima Foundation (to Y.S.), the Exploratory Research Center on Life and Living Systems (ExCELLS) (23EX601 and 25EX602 to Y.S.), and RIKEN (Pioneering Project to S.I. and Y.S.; RIKEN TRIP initiative “TRIP-AGIS” to S.I.). E.M.T. was a recipient of a JSPS Postdoctoral Fellowship for Research in Japan.

## Author contributions

Conceptualization, E.M.T. and S.I.;

Methodology, E.M.T., M.M., Y.S., and S.I.;

Formal analysis, E.M.T.;

Investigation, E.M.T.;

Writing – Original Draft, E.M.T. and S.I.;

Writing – Review & Editing, E.M.T., M.M., Y.S., and S.I.;

Visualization, E.M.T.;

Supervision, Y.S. and S.I.;

Project administration, S.I.;

Funding Acquisition, E.M.T., Y.S., and S.I.

---

**Figure S1.**
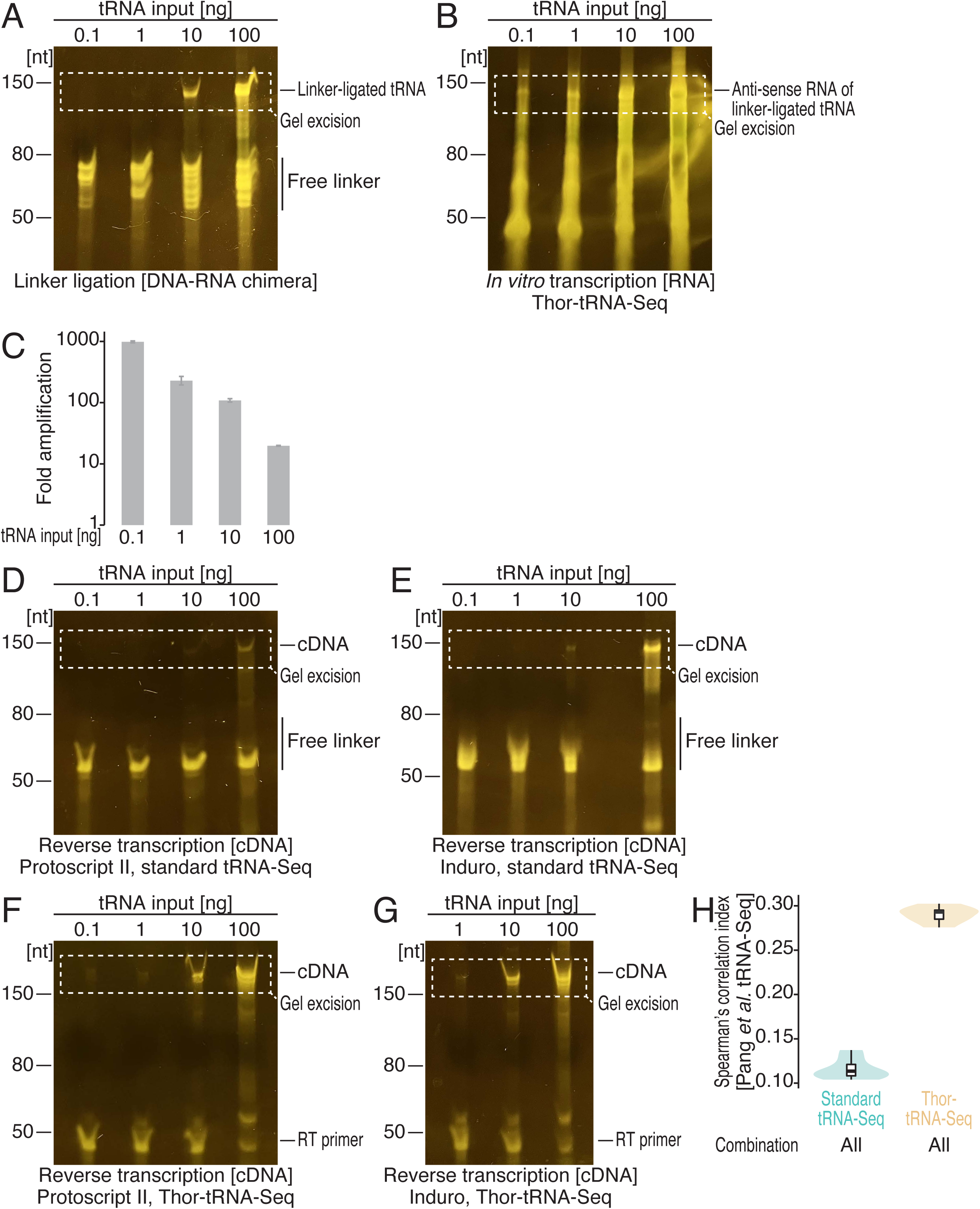
Characterization of Thor-tRNA-Seq library preparation, related to Figure 1. (A) Denaturing PAGE analysis of linker-ligated tRNAs across different inputs. Nucleic acids were stained with SYBR Gold. (B) Denaturing PAGE analysis of *in vitro* transcribed RNAs corresponding to tRNA antisense RNAs across different inputs. Nucleic acids were stained with SYBR Gold. (C) Fold amplification of RNAs from input tRNAs after *in vitro* transcription for the indicated conditions. The data are presented as the means (bars) for replicates (points, n = 2). (D and E) Denaturing PAGE analysis of reverse-transcribed cDNAs across different inputs. Protoscript II (D) and Induro (E) were used for the reverse transcription in standard tRNA-Seq. Nucleic acids were stained with SYBR Gold. (F and G) Denaturing PAGE analysis of reverse-transcribed cDNAs across different inputs. Protoscript II (D) and Induro (E) were used for the reverse transcription in Thor-tRNA-Seq. Nucleic acids were stained with SYBR Gold. (H) Box and violin plots of correlation levels comparing published tRNA-Seq data ^37^ with our data obtained in standard tRNA-Seq and Thor-tRNA-Seq.

## References

1. Gao, L., Behrens, A., Rodschinka, G., Forcelloni, S., Wani, S., Strasser, K., and Nedialkova, D.D. (2024). Selective gene expression maintains human tRNA anticodon pools during differentiation. Nat. Cell Biol. 26, 100–112.

2. Rak, R., Polonsky, M., Eizenberg-Magar, I., Mo, Y., Sakaguchi, Y., Mizrahi, O., Nachshon, A., Reich-Zeliger, S., Stern-Ginossar, N., Dahan, O., et al. (2021). Dynamic changes in tRNA modifications and abundance during T cell activation. Proc. Natl. Acad. Sci. U. S. A. 118, e2106556118.

3. Wilusz, J.E. (2015). Controlling translation via modulation of tRNA levels. Wiley Interdiscip. Rev. RNA 6, 453–470.

4. Kirchner, S., and Ignatova, Z. (2015). Emerging roles of tRNA in adaptive translation, signalling dynamics and disease. Nat. Rev. Genet. 16, 98–112.

5. Zhang, Z., Ye, Y., Gong, J., Ruan, H., Liu, C.-J., Xiang, Y., Cai, C., Guo, A.-Y., Ling, J., Diao, L., et al. (2018). Global analysis of tRNA and translation factor expression reveals a dynamic landscape of translational regulation in human cancers. Commun. Biol. 1, 1–11.

6. Dana, A., and Tuller, T. (2012). Determinants of Translation Elongation Speed and Ribosomal Profiling Biases in Mouse Embryonic Stem Cells. PLoS Comput. Biol. 8, e1002755.

7. Davyt, M., Bharti, N., and Ignatova, Z. (2023). Effect of mRNA/tRNA mutations on translation speed: Implications for human diseases. J. Biol. Chem. 299. 10.1016/j.jbc.2023.105089.

8. Sørensen, M.A., Kurland, C.G., and Pedersen, S. (1989). Codon usage determines translation rate in Escherichia coli. J. Mol. Biol. 207, 365–377.

9. Varenne, S., Buc, J., Lloubes, R., and Lazdunski, C. (1984). Translation is a non-uniform process. Effect of tRNA availability on the rate of elongation of nascent polypeptide chains. J. Mol. Biol. 180, 549–576.

10. Yu, C.-H., Dang, Y., Zhou, Z., Wu, C., Zhao, F., Sachs, M.S., and Liu, Y. (2015). Codon Usage Influences the Local Rate of Translation Elongation to Regulate Co-translational Protein Folding. Mol. Cell 59, 744–754.

11. Subramaniam, A.R., Zid, B.M., and O’Shea, E.K. (2014). An integrated approach reveals regulatory controls on bacterial translation elongation. Cell 159, 1200– 1211.

12. dos Reis, M., Savva, R., and Wernisch, L. (2004). Solving the riddle of codon usage preferences: a test for translational selection. Nucleic Acids Res. 32, 5036– 5044.

13. Parvathy, S.T., Udayasuriyan, V., and Bhadana, V. (2022). Codon usage bias. Mol. Biol. Rep. 49, 539–565.

14. Orellana, E.A., Siegal, E., and Gregory, R.I. (2022). tRNA dysregulation and disease. Nat. Rev. Genet. 23, 651–664.

15. Huang, S.-Q., Sun, B., Xiong, Z.-P., Shu, Y., Zhou, H.-H., Zhang, W., Xiong, J., and Li, Q. (2018). The dysregulation of tRNAs and tRNA derivatives in cancer. J. Exp. Clin. Cancer Res. 37, 101.

16. Yuan, W., Zhang, R., Lyu, H., Xiao, S., Guo, D., Zhang, Q., Ali, D.W., Michalak, M., Chen, X.-Z., Zhou, C., et al. (2024). Dysregulation of tRNA methylation in cancer: Mechanisms and targeting therapeutic strategies. Cell Death Discov. 10, 1– 19.

17. Padhiar, N.H., Katneni, U., Komar, A.A., Motorin, Y., and Kimchi-Sarfaty, C. (2024). Advances in methods for tRNA sequencing and quantification. Trends Genet. 40, 276–290.

18. Smith, T., Monti, M., Willis, A.E., and Kalmár, L. (2024). Benchmarking tRNA-Seq quantification approaches by realistic tRNA-Seq data simulation identifies two novel approaches with higher accuracy. eLife 13. 10.7554/eLife.96955.2.

19. Suzuki, T. (2021). The expanding world of tRNA modifications and their disease relevance. Nat. Rev. Mol. Cell Biol. 22, 375–392.

20. Marlow, K., and Su, Z. (2026). Hidden in plain sight: illuminating the tRNA landscape by sequencing. Genome Biol. 10.1186/s13059-026-03995-2.

21. Linderman, G.C., Zhao, J., Roulis, M., Bielecki, P., Flavell, R.A., Nadler, B., and Kluger, Y. (2022). Zero-preserving imputation of single-cell RNA-seq data. Nat. Commun. 13, 192.

22. Jiang, R., Sun, T., Song, D., and Li, J.J. (2022). Statistics or biology: the zero-inflation controversy about scRNA-seq data. Genome Biol. 23, 31.

23. Fazal, F.M., Han, S., Parker, K.R., Kaewsapsak, P., Xu, J., Boettiger, A.N., Chang, H.Y., and Ting, A.Y. (2019). Atlas of Subcellular RNA Localization Revealed by APEX-Seq. Cell 178, 473–490.e26.

24. Nair, R.R., Zabezhinsky, D., Gelin-Licht, R., Haas, B.J., Dyhr, M.C.A., Sperber, H.S., Nusbaum, C., and Gerst, J.E. (2021). Multiplexed mRNA assembly into ribonucleoprotein particles plays an operon-like role in the control of yeast cell physiology. Elife 10, e66050.

25. Sharma, E., Sterne-Weiler, T., O’Hanlon, D., and Blencowe, B.J. (2016). Global mapping of human RNA-RNA interactions. Mol. Cell 62, 618–626.

26. Cazenave, C., and Uhlenbeck, O.C. (1994). RNA template-directed RNA synthesis by T7 RNA polymerase. Proc. Natl. Acad. Sci. U. S. A. 91, 6972–6976.

27. Shichino, Y., Mito, M., Kinugasa, Y., Wakigawa, T., Yamashita, A., Mishima, Y., Imai, Y., and Iwasaki, S. (2025). Ultra-parallel ribosome profiling platform with RNA-dependent RNA amplification. bioRxiv, 2025.08.06.668979.

28. Milligan, J.F., Groebe, D.R., Witherell, G.W., and Uhlenbeck, O.C. (1987). Oligoribonucleotide synthesis using T7 RNA polymerase and synthetic DNA templates. Nucleic Acids Res. 15, 8783–8798.

29. Arnaud-Barbe, N., Cheynet-Sauvion, V., Oriol, G., Mandrand, B., and Mallet, F. (1998). Transcription of RNA templates by T7 RNA polymerase. Nucleic Acids Res. 26, 3550–3554.

30. Hoeijmakers, W.A.M., Bártfai, R., Françoijs, K.-J., and Stunnenberg, H.G. (2011). Linear amplification for deep sequencing. Nat. Protoc. 6, 1026–1036.

31. Behrens, A., Rodschinka, G., and Nedialkova, D.D. (2021). High-resolution quantitative profiling of tRNA abundance and modification status in eukaryotes by mim-tRNAseq. Mol. Cell 81, 1802–1815.e7.

32. Nakano, Y., Gamper, H., McGuigan, H., Maharjan, S., Li, J., Sun, Z., Yigit, E., Grünberg, S., Krishnan, K., Li, N.-S., et al. (2025). Genome-wide profiling of tRNA modifications by Induro-tRNAseq reveals coordinated changes. Nat. Commun. 16, 1047.

33. Cozen, A.E., Quartley, E., Holmes, A.D., Hrabeta-Robinson, E., Phizicky, E.M., and Lowe, T.M. (2015). ARM-seq: AlkB-facilitated RNA methylation sequencing reveals a complex landscape of modified tRNA fragments. Nat. Methods 12, 879– 884.

34. Zheng, G., Qin, Y., Clark, W.C., Dai, Q., Yi, C., He, C., Lambowitz, A.M., and Pan, T. (2015). Efficient and quantitative high-throughput tRNA sequencing. Nat. Methods 12, 835–837.

35. Zhang, R., Mayer, L., Hikida, H., Shichino, Y., Mito, M., Willemsen, A., Iwasaki, S., and Ogata, H. (2026). A giant virus forms a specialized subcellular environment within its amoeba host for efficient translation. Nat. Microbiol. 11, 584–596.

36. Chan, P.P., and Lowe, T.M. (2016). GtRNAdb 2.0: an expanded database of transfer RNA genes identified in complete and draft genomes. Nucleic Acids Res. 44, D184–D189.

37. Pang, Y.L.J., Abo, R., Levine, S.S., and Dedon, P.C. (2014). Diverse cell stresses induce unique patterns of tRNA up- and down-regulation: tRNA-seq for quantifying changes in tRNA copy number. Nucleic Acids Res. 42, e170.

38. Mito, M., Mishima, Y., and Iwasaki, S. (2020). Protocol for Disome Profiling to Survey Ribosome Collision in Humans and Zebrafish. STAR Protoc. 1, 100168.

39. Chen, S., Zhou, Y., Chen, Y., and Gu, J. (2018). fastp: an ultra-fast all-in-one FASTQ preprocessor. Bioinformatics 34, i884–i890.

40. Smith, T.S., Heger, A., and Sudbery, I. (2017). UMI-tools: Modelling sequencing errors in Unique Molecular Identifiers to improve quantification accuracy. Genome Res., gr.209601.116.

41. Dobin, A., Davis, C.A., Schlesinger, F., Drenkow, J., Zaleski, C., Jha, S., Batut, P., Chaisson, M., and Gingeras, T.R. (2013). STAR: ultrafast universal RNA-seq aligner. Bioinformatics 29, 15–21.

42. Li, H., Handsaker, B., Wysoker, A., Fennell, T., Ruan, J., Homer, N., Marth, G., Abecasis, G., Durbin, R., and 1000 Genome Project Data Processing Subgroup (2009). The Sequence Alignment/Map format and SAMtools. Bioinformatics 25, 2078–2079.

43. Morgan, M., Pagès, H., Obenchain, V., and Hayden, N. (2025). Rsamtools: Binary alignment (BAM), FASTA, variant call (BCF), and tabix file import. 10.18129/B9.bioc.Rsamtools.

